# Distinct microbiome composition of floor and air dust distort richness estimation in university dormitories

**DOI:** 10.1101/855684

**Authors:** Yu Sun, Yanling Li, Qianqian Yuan, Zefei Zhang, Yiqun Deng, Dan Norbäck, Xin Zhang, Xi Fu

**Affiliations:** Guangdong Provincial Key Laboratory of Protein Function and Regulation in Agricultural Organisms, College of Life Sciences, South China Agricultural University, Guangzhou, Guangdong, 510642, PR China; Key Laboratory of Zoonosis of Ministry of Agriculture and Rural Affairs, South China Agricultural University, Guangzhou, Guangdong, 510642, PR China; Institute of Environmental Science, Shanxi University, Taiyuan, PR China; Occupational and Environmental Medicine, Dept. of Medical Science, University Hospital, Uppsala University, 75237 Uppsala, Sweden; School of Public Health, Sun Yat-sen University, Guangzhou, PR China

## Abstract

The number of culture-independent indoor microbiome study has increased remarkably in recent years, but microbial composition among different sampling strategies remains poorly characterized and their impact to downstream microbiome analysis is also not clear. In this study, we reported a case study of microbial composition of floor and air dust in 87 dormitory rooms of Shanxi University, China. Floor and air dust were collected by vacuum cleaner and petri-dish, respectively, and the bacterial composition was characterized by 16S rRNA sequencing. The composition of floor and air dust differed significantly (R^2^ = 0.65, p < 0.001, Adonis), and *Pseudomonas* dominated in floor dust (75.1%) but was less common in air dust (1.9%). The top genera in air dust, including *Ralstonia* (15.6%), *Pelomonas* (11.3%), *Anoxybacillus* (9.3%) and *Ochrobactrum* (6.2%), all accounted for < 1% abundance in floor dust. The dominant *Pseudomonas* in floor dust swamped low frequency organisms, leading to significant lower number of operational taxonomic units (OTUs) compared with air dust in the same sequencing depth. The different microbial composition of floor and air dust can lead to differences in downstream bioinformatics analyses. We searched the dormitory microbiome against ~200,000 samples deposited in Microbiome Search Engine (MSE), and found that the compositions of floor dust samples were similar to samples from building environment and human nasopharynx, whereas the compositions of the air dust samples were similar to mosquito tissues.

**Importance:** Increasing number of indoor microbiome studies has been conducted in recent years, but the impact of sampling strategy is far from clear. In this study, we reported that the floor and air sampling can lead to drastic variation in microbial composition and downstream analyses in university dormitories, including microbial diversity estimation and compositional similarity search. Thus, the bioinformatics analysis results need to be interpreted with caution in microbiome studies, and it may be necessary to collected samples from different sampling strategies to comprehensively characterize the microbial composition and exposure to human occupants in an indoor environment.

## Introduction

The development and application of the cost-efficient next-generation sequencing technology in the past decade greatly facilitates the culture-independent researches to characterize microbiome composition in various ecological niches, including human gastrointestinal and respiratory tract, skin and indoor environment [1–6]. The first step of microbiome study is to collect samples, and various approaches has been applied for different sample types. The gut microbiome is commonly sampled from stool, which represents well the microbial composition of colonic lumen and mucosa [7], but other sampling strategies such as mucosal biopsy, brushing or rectal swab are also applied in various studies [8, 9]. Similarly, swab, spurum, lavage, aspirates and brushing have all been used in respiratory microbiome studies [10], and swab, biopsy, cup scrub, tape strip and surface scrape were used in skin microbiome studies [11]. It has been reported that sampling collection strategies produce detected variability in the gut microbiome studies. Thus it is necessary to understand the variations and standardize the laboratory and analysis protocols in the specific researches [12].

Compared to microbiome researches in human body sites, the number of studies in the built environment is much smaller and thus the variation between different sampling strategies is unclear. Several types of sampling strategies have been applied for the indoor studies, including floor or mattress dust collected by swab sampling or vacuum cleaner with special filters or socks, settled airborne dust collected by passive dust collectors such as petri dish and electrostatic dustfall collectors (EDCs) placed above the floor level, and airborne dust directly collected by air vacuum pump or BioSampler. The direct sampling of airborne dust is expected to be the most direct way to evaluate the inhalable exposure to indoor occupants, but the approach is relatively expansive, and since the microbial composition in air dust varies temporally, this approach represents short term microbial exposure. The settled dust represents long-term microbial exposure thus also possess unique strength in the indoor microbiome study. One study compared different sampling strategies and showed that the amount of microbial colonies and mycotoxins varied between floor vacuum dust and air dust [13]. However, this study used culture-dependent approach by counting the colony forming units, which did not present a comprehensive picture of overall microbial composition as this traditional approach can only culture less than 1% of total microbial species [14]. One recent study used culture-independent high-throughput sequencing approach to assess microbial composition of several passive dust collectors, such as petri dish, TefTex material and EDCs, and found these approaches collected similar microbial composition and quantity [15], but the widely used approach, vacuum settled dust, was not included and compared in the study. An indoor emission study on airborne allergenic fungi showed that 70% of indoor fungal aerosol particles were derived from indoor emission such as occupant resuspension, indicating similar fungal composition between air and floor dust [16].

In this study, we used culture-independent high-throughput technology to sequence amplicon gene in floor and settled airborne dust from 87 university dormitory rooms in 10 dormitory buildings in Shanxi University, and revealed highly distinct microbial composition from the two sampling approaches. Standard bioinformatics analyses were conducted and showed that the distinct microbial composition from different sampling strategies also led to drastic different downstream analysis results.

## Results

In this study, we used multiplexed Illumina high-throughput sequencing technique to characterize microbial composition in 87 randomly selected university dormitory rooms in ten buildings of Shanxi University, China. The settled floor dust was collected by vacuum cleaner for 4 minutes with a special cellulose acetate filter. The filter retains, and thus can capture the majority of the bacteria, which are typically in the range of 0.5-5 μm in length. Settled airborne dust was collected by petri-dish at a height of approximately 1.2m above the floor level for one week. Four floor dust and one air dust samples were used up in previous analysis or failed to amplified enough DNA for sequencing, thus in total 83 and 86 floor and air dust samples were sequenced. The gel electrophoresis of negative control showed that our samples were not contaminated by laboratory microbes (Figure S1). Chimeric and low-quality reads were filtered out before reads assembly and taxonomical classification. In total, 3,616 operational taxonomic units (OTUs) in 530 genera were identified.

### Distinct compositional difference between floor and airborne dust

We found drastic differences in bacterial composition between floor and airborne dust (Figure 1). Floor dust samples were dominated by Proteobacteria (89.3%, Figure 1A and Table S1), and the other Phyla only accounted for minority of the total bacterial load in university dormitories including Firmicutes (5.4%), Deinococcus-Thermus (3.0%) and Actinobacteria (1.1%). At the genus level, *Pseudomonas* accounted for 75.1% of total bacterial loads, followed by *Psychrobacter* (4.5%), *Cupriavidus* (4.2%), *Deinococcus* (2.6%), *Sphingomonas* (1.9%), *Bacillus* (1.8%) and *Planomicrobium* (1.4%, Figure 1B and Table S2). Most floor dust samples were dominated by *Pseudomonas* with a few exceptions; two samples from top floor of building No.9 had low abundance of *Pseudomonas* (<1%) but high abundance of *Deinococcus* (>30%) and *Cupriavidus* (>15%; Figure S2 and S3).

**Figure 1.**
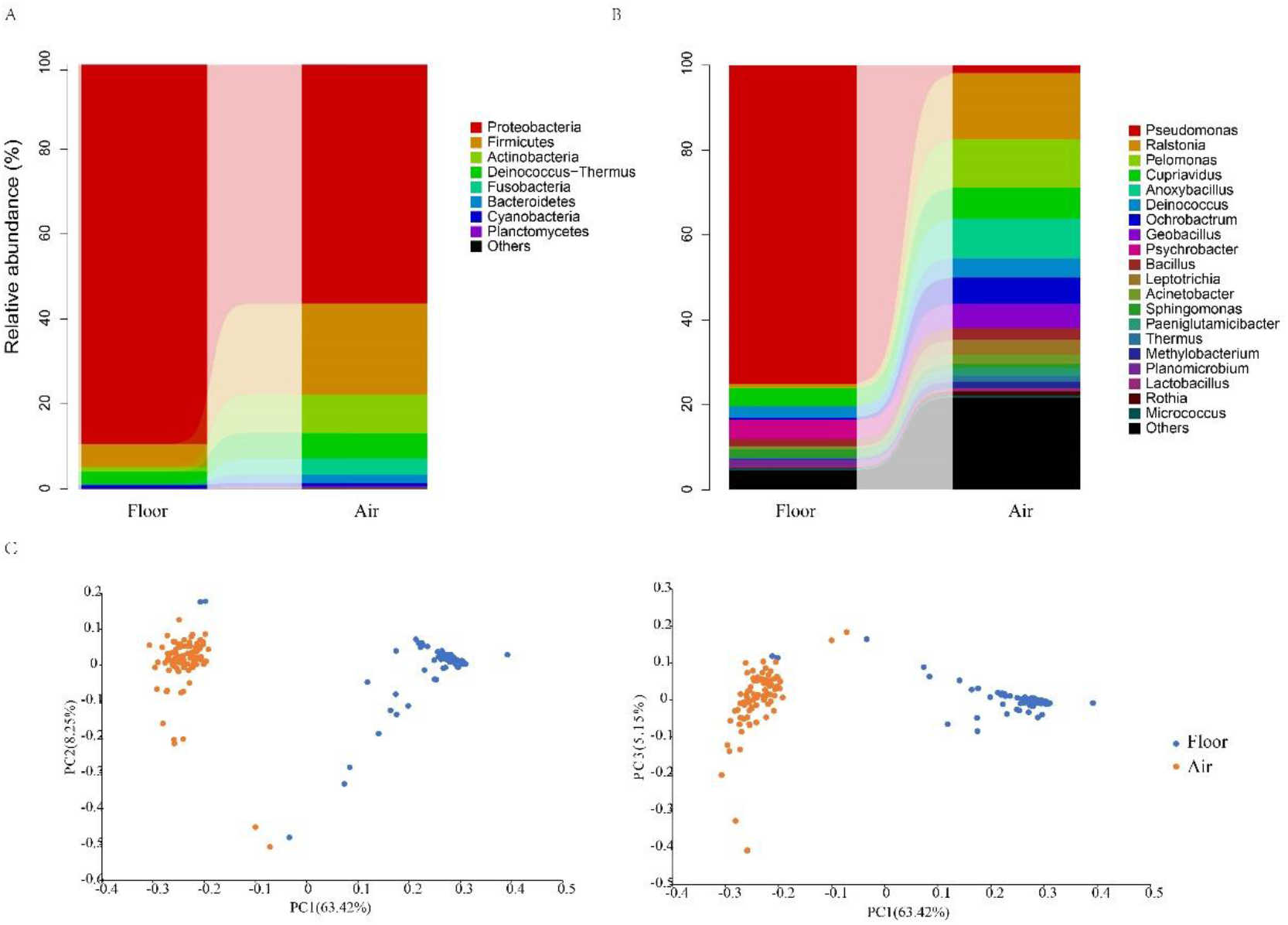
Bacterial relative abundance and community variation for floor and air dust. Relative abundance of bacterial taxa at phylum (A) and genus (B) level. (C) PCoA analyses of microbial composition based on weighted UniFrac distance was performed. Ordination plot between PC1 and PC2 and between PC1 and PC3 are displayed.

For the air dust, Proteobacteria accounted for 56.2% of all bacteria, followed by Firmicutes (21.5%), Actinobacteria (9.0%), Deinococcus-Thermus (6.0%), Fusobacteria (3.9%) and Bacteroidetes (2.0%, Figure 1A and Table S3). At the genus level, *Pseudomonas* was not as dominant in the air dust as in floor dust and only accounted for 1.9% of total bacteria (Figure 1B and Table S4). The top genera were *Ralstonia* (15.6%) and *Pelomonas* (11.3%), followed by *Anoxybacillus* (9.3%), *Cupriavidus* (7.3%), *Ochrobactrum* (6.2%), *Geobacillus* (5.7%), *Deinococcus* (4.6%), *Leptotrichia* (3.4%), *Bacillus* (2.7%) and *Acinetobacter* (2.3%). The microbial compositions were generally similar in all air dust except in two rooms of building No.1, which had more than 80% of *Bacillus* (Figure S3).

The differences in microbial compositions between floor and air dust were also confirmed by the quantitative test such as principle coordinate analysis (PCoA) and Adonis. Samples from the two sampling methods were separated along axis 1, and the axis dominantly explained 63.4% of total variation in the dataset (Figure 1C). The two floor dust samples from building No.9 were compositionally similar to the air dust. Adonis test showed similar result that sampling type explained 65% of total variance in the dataset (p < 0.001), whereas other environmental factors, such as building age and location, sex of occupants, wall surface type and cleaning frequency, were not significantly associated with total microbial variation (p > 0.05).

### Diversity estimation and composition similarity search in public database

We conducted richness analysis based on 20 repeated rarefaction analyses on even depth of 11,000 sequences for all sample. Floor dust had significant lower number of observed OTUs compared to the air dust (p < 0.001, Man-Whitney U test; floor dust 345.1 ± 82.3, air dust 906.2 ± 179.6, mean ± standard deviation). Similar result was obtained for richness estimation at the genus level (p < 0.001, floor dust 56.5 ± 22.0, air dust 184.0 ± 32.5). The species accumulation curve analysis showed that the two sampling methods had different accumulation pattern (Figure 2). The first 20 samples of air dust caught the majority of OTUs, whereas the number of accumulated OTUs was still increasing after 80 samples for floor dust. The results suggest that either the microbial OTUs differs remarkably within the floor dust samples, or the sequencing depth of floor dust is not deep enough to uncover certain low frequency bacteria in each sample.

**Figure 2.**
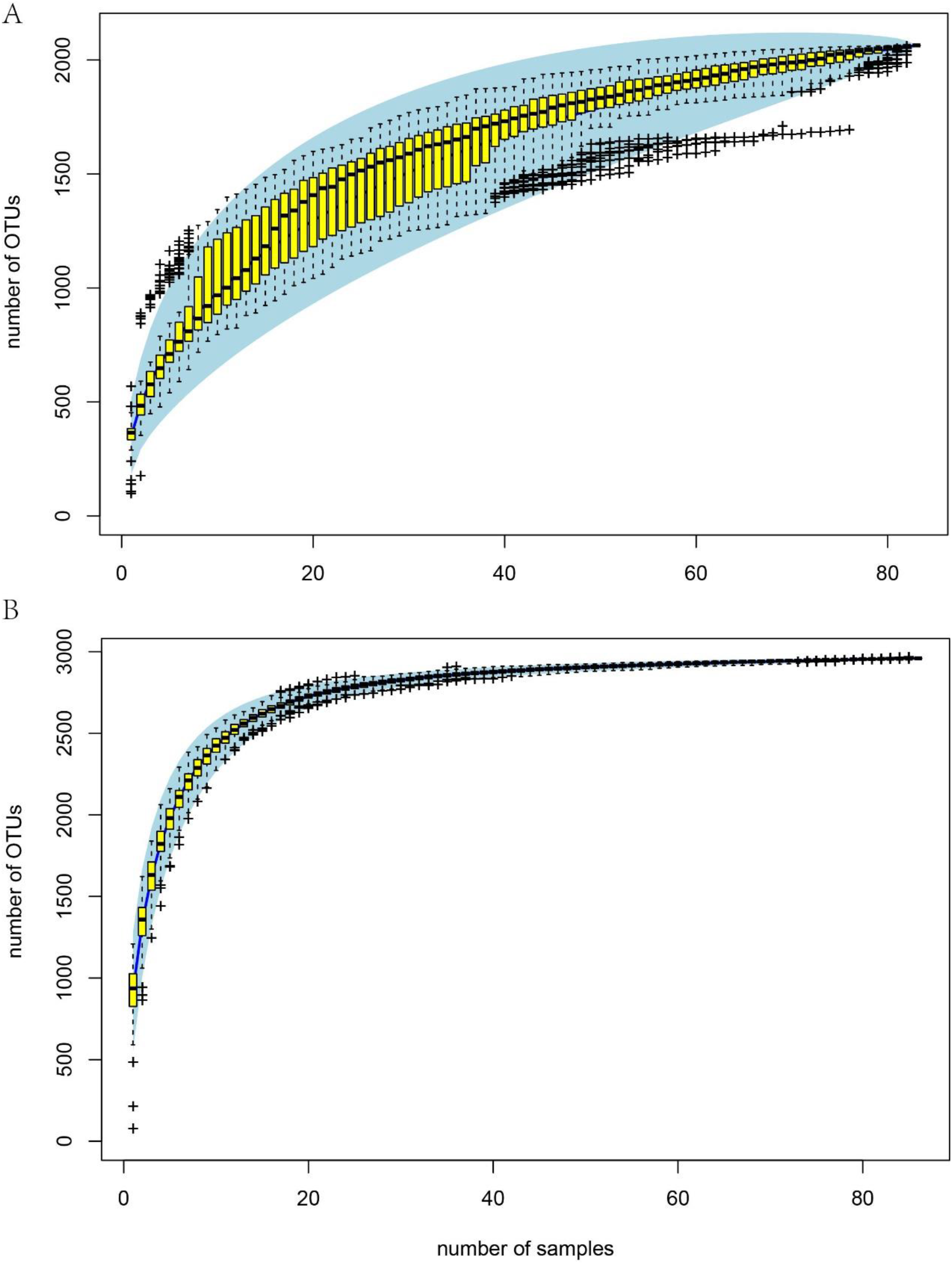
Species accumulation curve for (A) floor and (B) air dust. The dark black line is the median species accumulation curve from 100 random permutations of the data. The yellow box represents Q1 (25%) and Q3 (75%) quartile. The top whisker represents Q3 + 1.5*IQR (interquartile range) and the bottom whisker represents Q1 - 1.5*IQR, and the “+” symbol represents outlier in the analysis. The shaded light blue area represents 2 times the standard deviations.

The differences in microbial compositions between floor and air dust also lead to variation in the downstream bioinformatics analyses. Microbiome Search Engine (MSE) [17] is a recently developed online analysis platform that enables rapid search of query microbiome against more than 200,000 bacterial microbiome samples from Qiita [18]. The online platform can not only find the most compositionally similar microbiome samples in the dataset, but can also report a novelty score which is a quantitatively index assessing the novelty of the query microbiome. The low similarity between the query microbiome and the top ten best matched microbiome indicates the query microbiome is very “novel” and a high novelty score will be given to the query microbiome. We searched the floor and air dust samples in the MSE platform. For 2/3 of the floor dust samples, the most similar microbiome were from mosquito tissues, whereas the remaining samples were best matched with samples from birds, soil and buildings (Table 1 and S5). Approximately 60% of air dust samples from dormitory rooms were best matched with building environment and human nasopharynx (Table S6). The high similarity between dormitory floor dust and mosquito tissue microbiome was due to the high percentage of *Pseudomonas* in both sample types (Table S5). In addition, due to the high similarity between floor dust and mosquito microbiome, the novelty scores of the floor dust samples were significantly lower compared with the air dust samples (p < 0.001).

**Table 1.** Number and percentage of compositionally best matched samples identified in the Microbiome Search Engine (MSE) for the dormitory floor and air dust.

| Habitat of top best matched samples | Number of samples in floor dust (%) | Number of samples in air dust (total 86) |
| --- | --- | --- |
| Mosquito | 56 (67.5%) | 0 (0%) |
| Bird | 5 (6.0%) | 10 (11.6%) |
| Marine | 2 (2.4%) | 0 (0%) |
| Soil | 6 (7.2%) | 10 (11.6%) |
| Skin | 4 (4.8%) | 7 (8.1%) |
| Building | 3 (3.6%) | 32 (37.2%) |
| Nasopharynx | 0 (0%) | 19 (22.1%) |
| Others | 7 (8.4%) | 8 (9.3%) |

## Discussion

In this study, we conducted amplicon sequencing of 16S rRNA for floor dust and settled airborne dust in dormitory rooms of Shanxi University, China. We found drastic difference in microbial composition for the two sampling strategies. The sampling strategy explained two thirds of compositional variation, whereas all other environmental factors contributed minimally.

The negative control, which contains only reagent but no dust samples, showed no band in the gel electrophoresis. Also, all the floor and air dust samples were extracted, amplified and sequenced by the same protocol, and all sequencing was conducted in a single run of Miseq which eliminated the potential run-to-run variability. Thus the variation between the floor and air dust samples is not due to contamination or technical bias, but rather due to biological difference in the samples. It has been suggested that the microbial composition can differ between floor and airborne dust, although most of the claims were verified by the culture-dependent study [13]. The vacuum dust was reported to be inefficiently in capture small airborne microbes, leading to their underrepresentation compared with the large body size taxa [19, 20]. Floor dust contains outdoor dust or soil material that is brought in by shoes of the occupants. The Chinese dormitory room is generally occupied by 3 to 6 students and thus relatively higher amount of outdoor dust could be transferred to indoor rooms by the occupants. *Pseudomonas* is one of the most diversely distributed and ubiquitous bacterial genera in the outdoor environment, such as soil, water, sediments or plants and animals [21]. *Pseudomonas* brought in by shoes can attach to large soil particles which were seldom stirred up by occupants to the level higher than 1.2 m, the height of the petri-dish placed to collect airborne dust. Also, *Pseudomonas* was reported to be a large size rod-shape bacteria, with ~1.5-2.2 μm in diameter and up to 5.0 μm in length, much larger than some of the common indoor bacteria, such as *Staphylococcus* (~0.5-0.8 μm) and *Bacillus* (~0.5-0.8 μm) [22]. The large body size of *Pseudomonas* or the attachment of the microbes to large particles could lead to the accumulation on the floor surface and limited resuspension into the air, contributing to the compositional variation in dormitories. In addition, *Pseudomonas* has been found in tap water [23]. Mop cleaning with water is performed daily in all dormitories of this study, and can transfer *Pseudomonas* from tap water to the floor. Incomplete cleaning of mop could also lead to accumulation or even growth of *Pseudomonas* in the mop. It should be noted that the result reported in this study does not imply the drastic difference should be consistently found between floor and air dust in other indoor environment. One fungal particle emission study reported the 80% of airborne allergenic fungal taxa were derived from floor dust resuspension in classrooms of seven primary schools, indicating similar fungal composition between floor and air dust [16]. Thus, it may be more appropriate to interpret the results in the dormitory rooms of Shanxi University as an individual case. More studies are needed to build an overall picture of indoor microbiome composition from different sampling strategies.

We found significant higher microbial richness in air dust compared with floor dust in dormitories, and argue that the phenomenon is more likely due to technical artefact rather than biological fact. Intuitively, it is hard to imagine that the diversity of floor microbiome is only ~35% of airborne microbiome in the indoor environment, since the majority of microbes should eventually settle down to the ground if no force constantly stirs up and resuspends floor dust. Indeed, a few studies reported microbial richness in floor dust is generally higher than richness in air dust. One school study reported fungal richness in floor dust was 1.4 times the fungal richness in air dust [16], and a crawling infant study reported that the bacterial richness (estimated by Chao1 and Shannon diversity) in the floor carpet was approximately one fold higher than the bacterial richness in infant breathing air (25 cm above ground) and two fold higher than adult breathing air (1.5 m above ground) [24]. Current microbiome analysis standard generally set same rarefaction depth for all samples within a study, and in this study the depth was set to 11,000 reads. However, as 3/4 of reads in floor dust were *Pseudomonas*, there were only less than 3,000 reads derived from all other organisms, and certain proportion of low frequency microbes (<0.1%) may not have been sequenced with this depth. This has also been indicated by the accumulation analysis of Figure 2. A previous study showed that a mycology classroom had a significantly lower richness than the other nonmycology classrooms, due to the dominance effect of the sporulating fungal puffballs [25]. The puffballs accounted for ~70% of total fungal load, which is similar to the dominant abundance of *Pseudomonas* observed in the floor dust samples. The study suggested that the dominant distortion was associated with elevated biomass derived from the dominant organisms, and sequencing higher depth in the corresponding samples could solve the richness distortion. However, current microbiome protocol generally aim to sequence all samples in the same depth to ensure the comparability in the downstream analysis, and analyzing samples with various sequencing depth could lead to many other issues and biases. Even this is possible in certain situation, how many reads and depth should be added in the specific sample is another open question.

Except richness estimation, the distinct microbial composition of floor and air dust also lead to variation in bioinformatics analyses. Approximately 2/3 of the floor dust samples best matched to mosquito tissues sampled in southern Ontario, Canada [26]. Mosquitos are present in dormitory rooms especially in summer seasons, but due to the small biomass amount, they are unlikely to contribute majorly to the indoor microbiome composition. The reverse transmission route from indoor dust to mosquito surface can occur, but the high abundance of *Pseudomonas* in mosquitos are commonly found in the gastrointestinal tract rather than on the surfaces, and the *Pseudomonas* are essential for mosquito development, digestion, fecundity and even preventing *Wolbachia* infection [27–29]. Indeed, an in-depth OTU analysis with a 97% sequence similarity cutoff (approximately species level resolution) for one dust sample showed that *Pseudomonas* OTUs present in indoor dust was different from OTUs in mosquito gut [30, 31]. The high similarity between floor dust and mosquitos microbiome indicates that the current amplicon sequencing and analysis pipeline are generally optimized at genus or above taxonomic level, and may have limitations to distinguish sensitive variation at the species and strain levels. Thus, future studies with more accurate taxonomic resolution, such as shotgun metagenomics or full-length amplicon sequencing by third-generation instrument, are needed to resolve the ambiguity in microbiome composition. Compared to the floor dust samples, ~60% of air dust samples best matched compositional with building environment and human nasopharynx samples, which is more in line with the intuitive expectation. Thus, caution is needed in interpreting microbiome data and analysis results even for samples from the same room but with different sampling strategies.

## Methods

Floor dust and air born dust were collected from 87 rooms in 10 dormitory buildings in Shanxi University, Taiyuan, China, 8-10 rooms in each building. Some samples failed for the DNA failed for the DNA quality test, and in total 83 floor and 86 air dust samples were qualified for DNA extraction and amplification for sequencing.

### Dust sample collection

Floor dust was collected by a 400 W vacuum cleaner with a dust sampling device (ALK Abello, Copenhagen, Denmark) equipped with a Milipore filter (pore size 6 μm). The filter is made of cellulose acetate and, according to manufacturer’s instruction, the filter retains 74% of particles 0.3-0.5 μm, 81% of particles 0.5-1.0 μm, 95% of particles 1-10 μm and 100% of larger particles (> 10 μm) [32]. The vacuum sampling procedure lasted for 4 minutes, including 2 minutes on floor and 2 minutes on other surfaces like beds and desks. The vacuum procedure was conducted twice at both window and corridor side in each dormitory room, and the two floor dust samples were merged for sequencing. Airborne settled dust was collected by two opened petri-dishes laying on a flat surface around 1.2 miters high for seven days. Next step in the lab, the floor dust samples were sieved to fine dust through a 0.3-mm mesh screen, and suspended in PBST buffer. The petri-dishes were washed by 2ml PBST buffer and stored in Eppendorf tubes. Both of the supernatant from floor dust and air dust were stored in the freezer at −80°C and later used for DNA extraction.

### Bacterial DNA extraction and amplicon sequencing

Total genomic DNA were extracted by E.Z.N.A. Soil DNA Kit D5625-01 (Omega Bio-Tek, Inc., Norcross, GA, USA), followed manufacturer’s instruction. Bead beating and spin filter technique were used for DNA extraction. Extracted DNA quantity and quality were evaluated by NanoDrop ND-1000 spectrophotometer, agarose gel electrophoresis and Microplate reader (BioTek, FLx800). Negative control with only reagent was added to evaluate laboratory microbial contaminations in the agarose gel electrophoresis. Forward primer 338F (ACTCCTACGGGAGGCAGCA) and reverse primer 806R (GGACTACHVGGGTWTCTAAT) were chosen for the 16s rRNA gene V3V4 region amplification. Sample specified 7-bp barcode sequences were incorporated into the multiplexing step. PCR reagents included 1 μl (10 μM) of forward and reverse primers, 2 μl DNA template, 5 μl Q5 reaction buffer, 2 μl (2.5 mM) dNTPs, 5 μl Q5 High-Fidelity GC buffer, 0.25 μl Q5 High-Fidelity DNA polymerase and 8.75 μl of ddH_2_O. PCR started with 2 min 98 °C denaturation, followed by 25 cycles of 15s 98 °Cdenaturation, 30s 55 °C annealing and 30s 72 °C extension process, and the whole PCR process ended with a 5 min extension at 72 °C. PCR amplicons were purified by Agencourt Beads (Bechman Coulter, Indianapolis, USA) and quantified by PicoGreen dsDNA Assay Kit (Invitrogen, CA, USA). In total, 169 samples were successfully amplified and further sequenced by Illumina MiSeq platform with MiSeq Reagent Kit v3. DNA extraction and multiplexed high-throughput sequencing were conducted by Personalbio (www.personalbio.cn).

### Bioinformatics and microbiome analysis

Raw sequences were extracted according to the barcode information and filtered with following criteria: minimum reads sequence length > 150 bp, average Phred score >20, contained no ambiguous bases and no mononucleotide repeats that > 8 bp [33]. Chimeric reads were removed by USEARCH (v5.2.236) [34], and paired-end reads were assembled by FLASH v(1.2.7) [35], with minimum 10 bp overlapping between forward and reverse reads without mismatches. The assembled high-quality reads were clustered into operational taxonomic units (OTUs) at 97% sequence similarity threshold by UCLUST [34]. A representative sequence from each OTUs was selected to blast against the Silva ribosomal RNA database [36] to acquire the taxonomic information by using the best hit. The following analyses were conducted in Quantitative Insights Into Microbial Ecology (QIIME, v1.8.0) platform [37] and R. An OTU table was built to store the abundance for each taxonomic units. The operational taxonomic unit threshold (c value) was set to 0.01% in QIIME with other parameters following a previous suggestion [38]. All samples were rarefied to even depth of 11,000 reads for the diversity analyses. Beta diversity analysis were conducted by using weighted UniFrac distance metrics [39], and visualized by principle co-ordinate analysis (PCoA) [40]. The microbiome similarity search was conducted on the Microbiome Search Engine (MSE, http://mse.single-cell.cn/)[17], and the query samples were first pre-processed by Parallel-META 3 [41, 42].

## Declarations

### Ethics approval and consent to participate

Not applicable.

### Consent for publication

Not applicable.

### Availability of data and material

Sequencing data was deposited in Qiita with study ID 12841 (https://qiita.ucsd.edu/study/description/12841).

### Authors’ contributions

ZZ and XZ collected dust samples in hotels. YS and XF designed the project. YS, YL, QY and XF carried out the analysis. YS, DN, XZ and XF drafted the manuscript. All authors read and approved the final manuscript.

## Acknowledgments

We thank Personalbio (www.personalbio.cn) for help in sequencing analysis.

## Supplementary Figures

**Figure S1.**
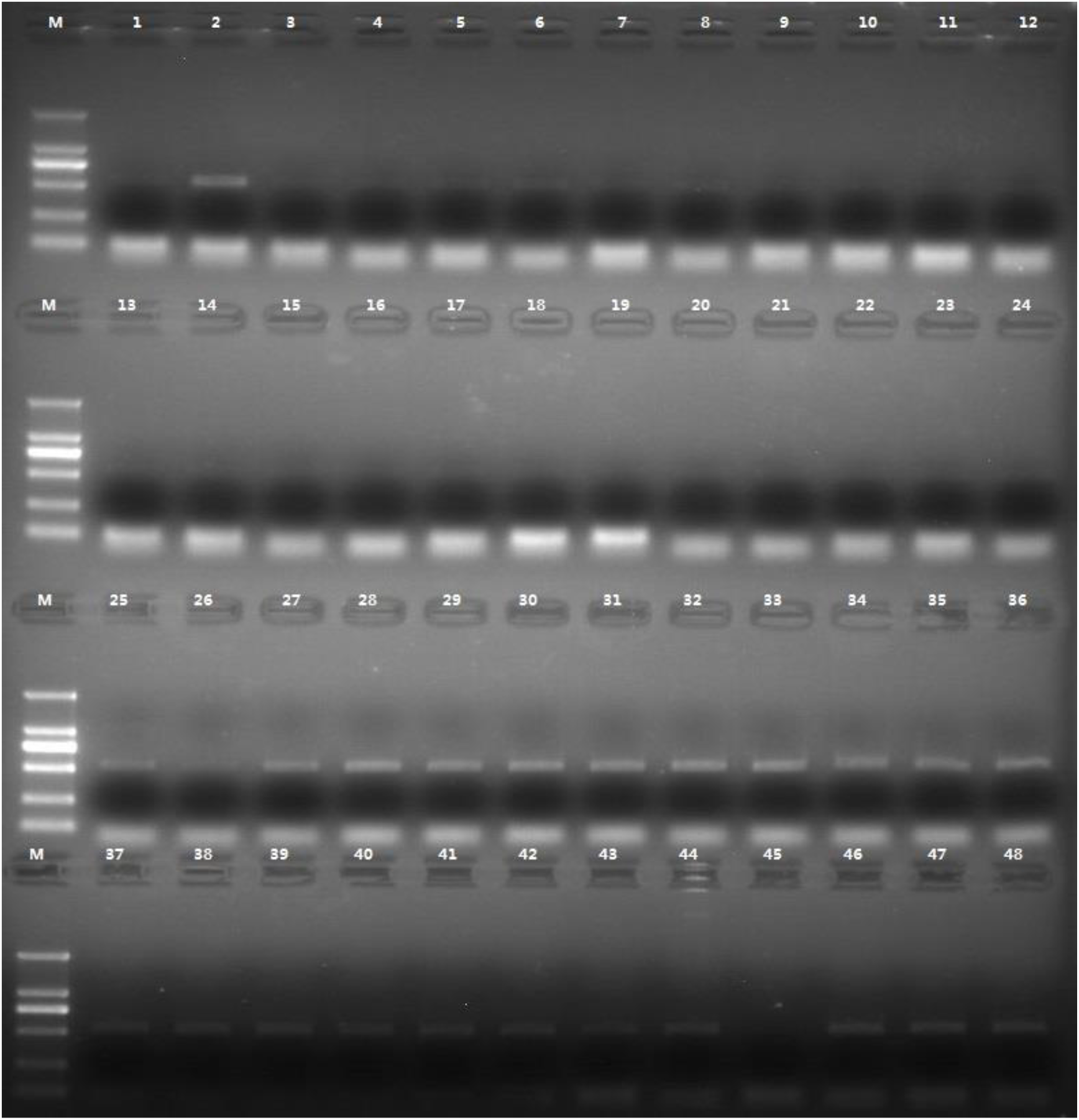
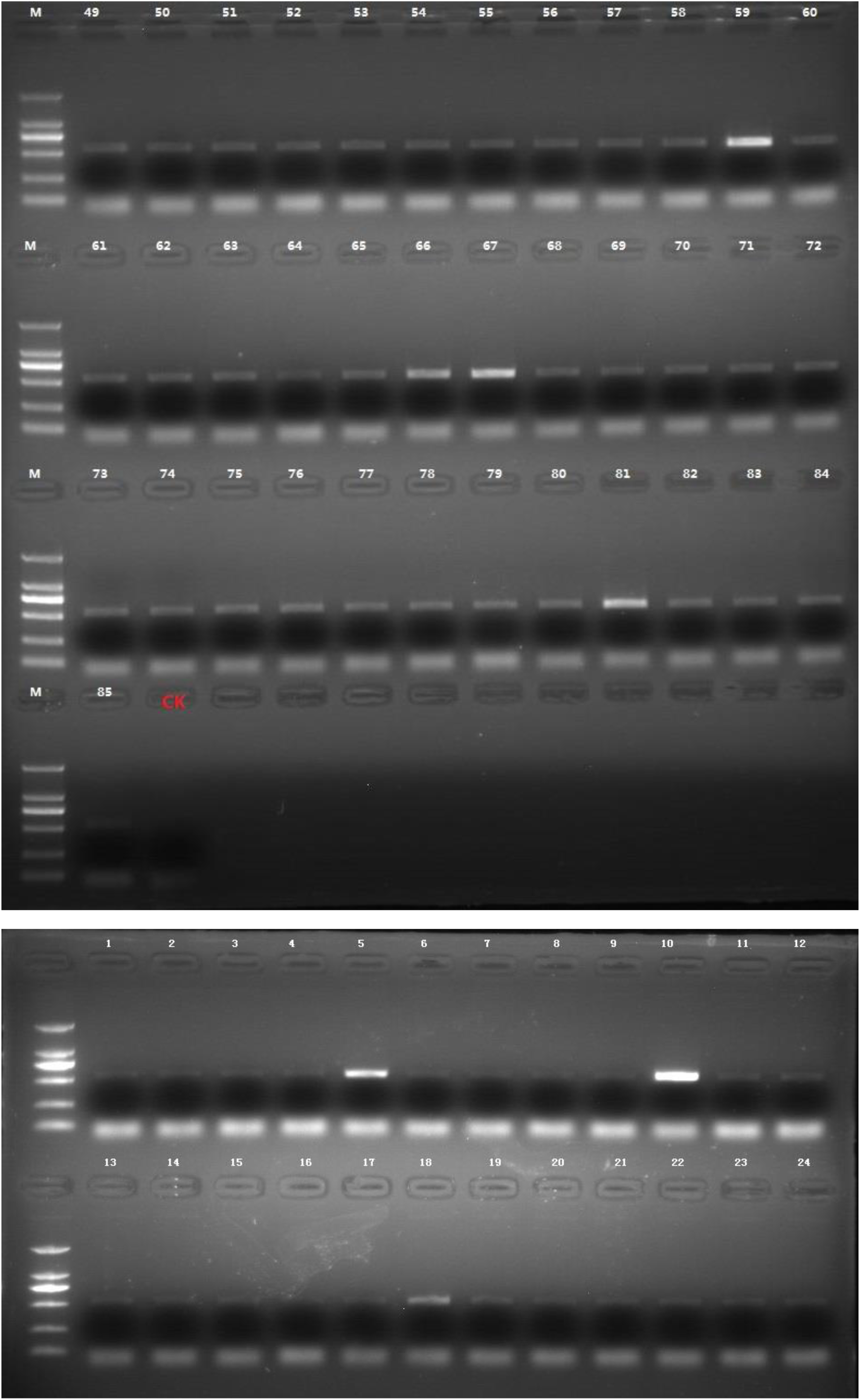
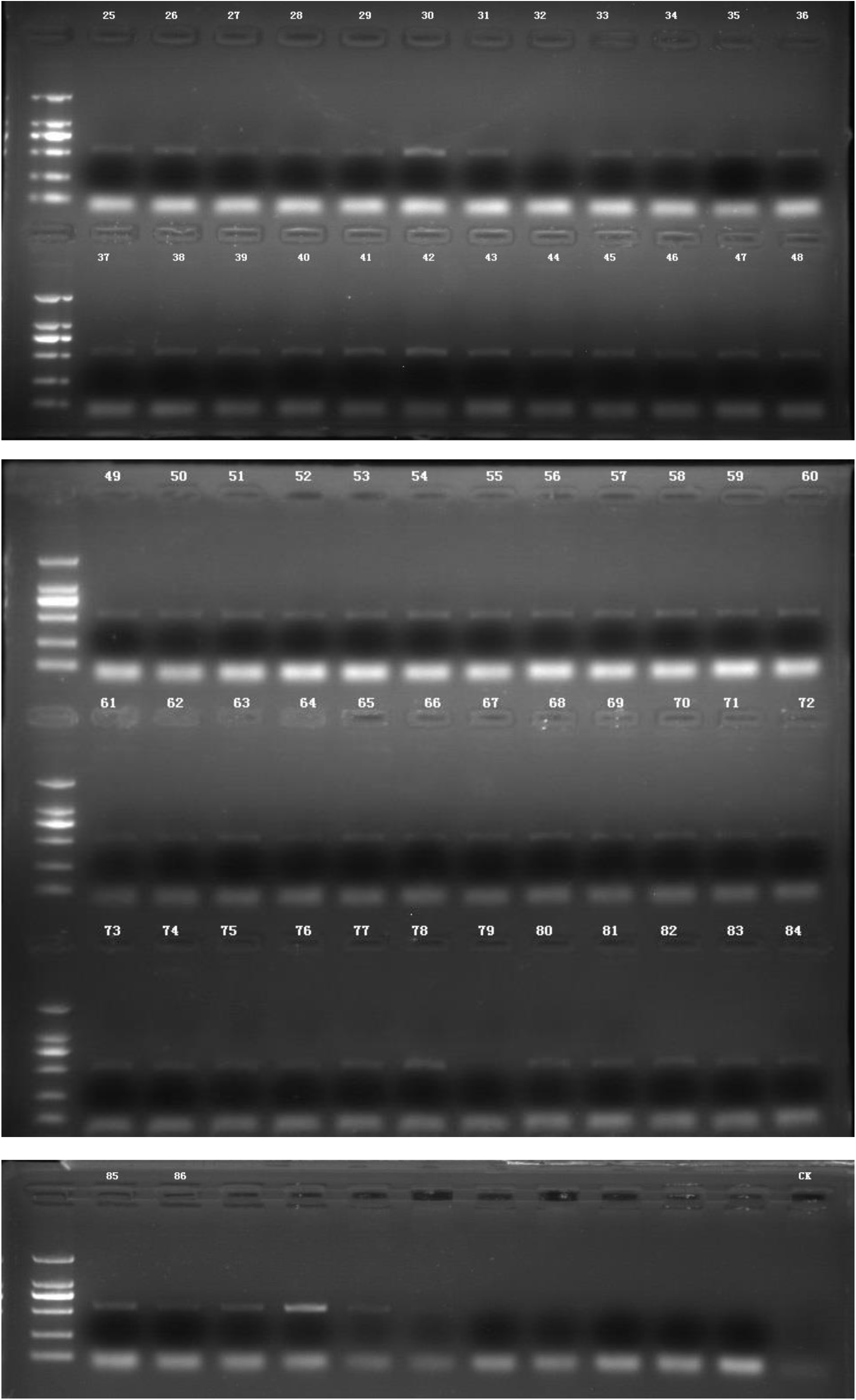
Gel electrophoresis of amplified 16S rRNA in the floor (top 1-85) and the air dust samples (bottom 1-86) and a negative control marked as CK.

**Figure S2.**
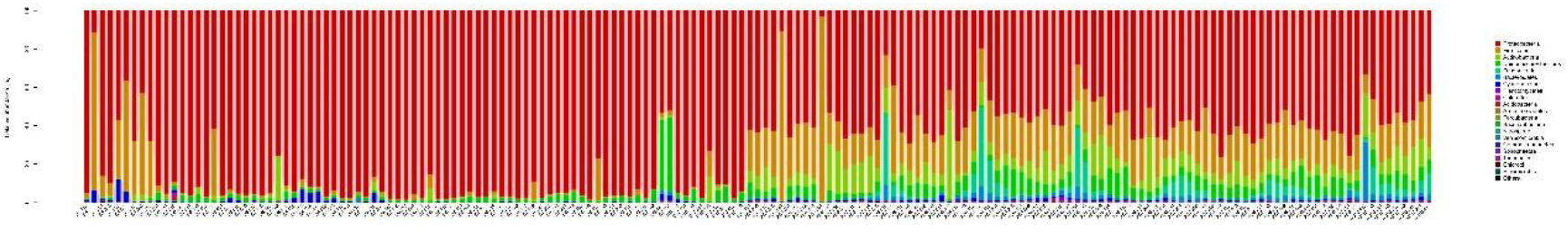
Bacterial relative abundance at the phylum level for all floor (left) and air dust samples (right).

**Figure S3.**
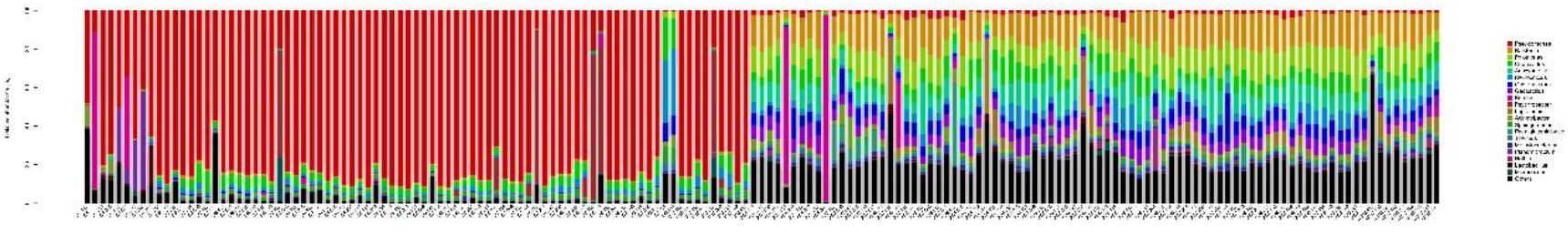
Bacterial relative abundance at the genus level for all floor (left) and air dust samples (right).

